# A Graph Theoretical Approach to Data Fusion

**DOI:** 10.1101/025262

**Authors:** Justina Žurauskienė, Paul DW Kirk, Michael PH Stumpf

## Abstract

The rapid development of high throughput experimental techniques has resulted in a growing diversity of genomic datasets being produced and requiring analysis. A variety of computational techniques allow us to analyse such data and to model the biological processes behind them. However, it is increasingly being recognised that we can gain deeper understanding by combining the insights obtained from multiple, diverse datasets. We therefore require scalable computational approaches for data fusion.

We propose a novel methodology for scalable unsupervised data fusion. Our technique exploits network representations of the data in order to identify (and quantify) similarities among the datasets. We may work within the Bayesian formalism, using Bayesian nonparametric approaches to model each dataset; or (for fast, approximate, and massive scale data fusion) can naturally switch to more heuristic modelling techniques. An advantage of the proposed approach is that each dataset can initially be modelled independently (and therefore in parallel), before applying a fast post-processing step in order to perform data fusion. This allows us to incorporate new experimental data in an online fashion, without having to rerun all of the analysis. The methodology can be applied to genomic scale datasets and we demonstrate its applicability on examples from the literature, using a broad range of genomic datasets, and also on a recent gene expression dataset from *Sporadic inclusion body myositis Availability*. Example R code and instructions are available from https://sites.google.com/site/gtadatafusion/.

## Introduction

Given the broad variety of high-throughput biological datasets now being generated, attention is increasingly focusing on methods for their integration.^1–3^ Different technologies allow us to probe different aspects of biological systems, and combining these complementary perspectives can yield greater insight.^4,5^

Methods for data fusion are rapidly evolving, and moving towards potential clinical applications.^4^ For example, recently proposed methods try to identify cancer subtypes using fused similarity networks applied to a combination of DNA methylation, mRNA expression and miRNA expression datasets.^3^ Cancer subtype discovery has also been a focus of other recent data integration efforts.^2,5,6^

Data integration methods can be categorised as either unsupervised (the subject of the present work) or supervised^7–10^. Existing unsupervised techniques for the fusion of multiple (i.e. more than two) datasets fall into one of two broad categories: those which are Bayesian^1,2^ and those which are not.^3,11^ For many real-world or large-scale analyses the lack of suitable training sets places severe limitations on supervised algorithms, and unsupervised learning methods have thus gained in popularity. Current Bayesian approaches for unsupervised data fusion rely on mixture model representations of each dataset, with dependencies between datasets modelled either using coefficients that describe the similarity between pairs of datasets,^1^ or by assuming that each dataset has a structure that adheres – to a lesser or greater degree – to an overall consensus structure that is common to all datasets.^2^ Bayesian approaches have the advantage of having firm probabilistic foundations, of forcing all prior beliefs and assumptions to be formally expressed at the start of the analysis, and of allowing the uncertainty in unknown quantities to be encapsulated in (samples from) posterior densities. For these reasons, Bayesian approaches are usually to be preferred; however, computational cost may prohibit their application to large (e.g. genome-scale) datasets.

In this work we introduce a new approach for the fusion of heterogeneous datasets. Our methodology is closely related to the graph-theoretic approach, which may be used to test for associations between disparate sources of data.^12^ Our approach has two basic steps. In the first, we obtain (independently) for each dataset either an ensemble of networks (in the Bayesian case), or a single network (in the non-Bayesian case), where networks are used to represent the structure and dependencies present in the dataset. In the second, we perform a post-processing step which compares the networks obtained for different datasets, in order to identify common edges, and to thereby assess and quantify the similarities in dataset structure.

Although, in principle, we could consider any type of structure that may be represented by a network, in the present work we are specifically interested in identifying and comparing the clustering structure possessed by each dataset, and it is at this level that we perform data fusion (Figure 1). In the Bayesian case, we employ for the purpose of cluster identification Dirichlet process mixture (DPM) models with either Gaussian process (GP) or multinomial likelihood functions. In our approach, data fusion is performed by constructing the connectivity networks that represent each clustering, and then forming “consensus networks” that identify the clusters upon which multiple datasets agree.

**Figure 1.**
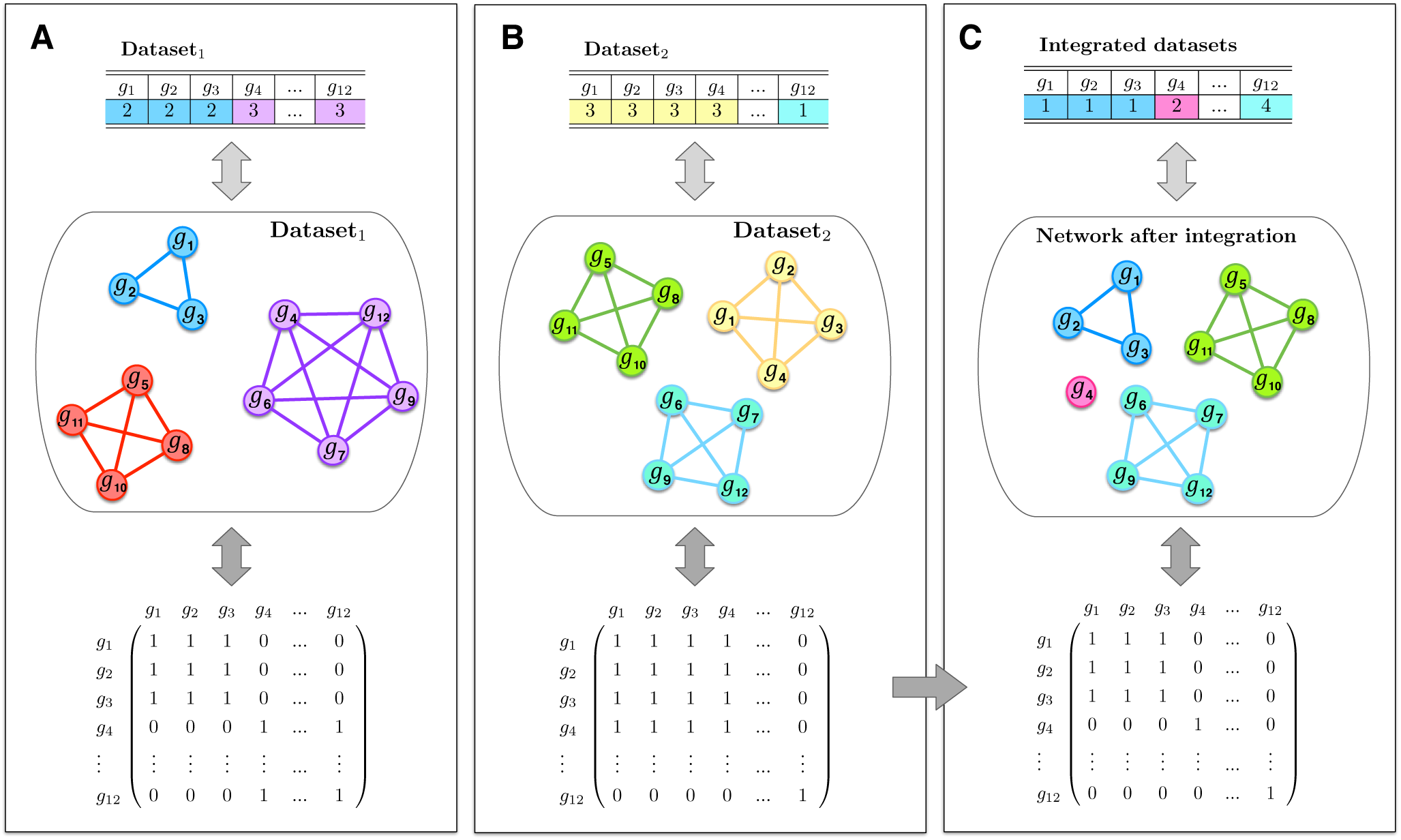
Method illustration. It is common to visualise the clustering outcome in a “table-like” fashion, by listing all genes next to their associated cluster labels. To visualise this, we construct corresponding graphs; here, each node in the network represents a gene and a line indicates that two genes cluster together. We use different colour schemes to represent cluster labels. By adopting a graph-theoretical approach we can represent each network as an adjacency matrix, which in turn can be used for data integration. **(A)** Illustrates an artificial example of 12 genes that are assigned into three clusters (e.g. control case) from the first dataset. **(B)** Illustrates the same list of genes assigned into different three clusters (e.g. disease case) from a second dataset. **(C)** Illustrates the corresponding network (and cluster assignment) after performing data integration step.

The integration step is somewhat similar to the consensus clustering approach,^13^ which was developed for the purpose of assessing cluster stability and for identifying the consensus across multiple evaluations of the same clustering approach. In contrast with other existing techniques,^1,2^ in our approach each dataset is initially considered independently. Although this is likely to result in some loss of sensitivity, it also means that computations can be performed in a parallel fashion, which offers potentially significant advantages in cases where we are dealing with large datasets, or where it is necessary to rerun computations in order to consider additional datasets.

## Results

Here we develop a novel *graph theoretical approach* for integrating clustering outcomes across several datasets, which we refer to as “GTA” for brevity. GTA may be applied to the output of Bayesian or non-Bayesian clustering algorithms.

### Data integration

There are many different methods for data integration, most of which set out to accomplish one (or both) of the following two key aims: (i) modelling the dependencies that exist within and between datasets; and (ii) combining the predictions derived from one dataset with those derived from another. Assuming for convenience of notation that we wish to integrate just a pair of datasets, we might consider that the “ideal” way to fulfil the first of these two aims would be to model the joint distribution, *p*(*q*^(1)^, *q*^(2)^ | *D*^(1)^, *D*^(2)^), of the predictions, *q*^(1)^ and *q*^(2)^, derived from datasets, *D*^(1)^ and *D*^(2)^ (respectively)^1^. Such an approach poses many potential challenges, not least that the datasets may be of very different types and/or may be of (very) high dimension. In contrast, the second aim just requires us to define a *fusion function*, *f*, for combining the predictions. We shall assume that this function is deterministic; i.e. for given predictions, *q*^(1)^ and *q*^(2)^, we assume that *f* maps the predictions onto a single combined output, *f* (*q*^(1)^, *q*^(2)^). One challenge in this case is to assess the uncertainty/confidence in this output. If we were able to sample *M* pairs, 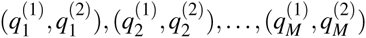 from the joint distribution, *p*(*q*^(1)^, *q*^(2)^ | *D*^(1)^, *D*^(2)^), then we could consider the set 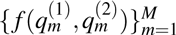 in order to assess the variability in the output.

In general, modelling the joint distribution, *p*(*q*^(1)^, *q*^(2)^ *D*^(1)^, *D*^(2)^), is much more computationally demanding than defining a fusion function. To make modelling *p*(*q*^(1)^, *q*^(2)^ | *D*^(1)^, *D*^(2)^) more tractable, here we make an independence assumption and factorise the joint distribution as,

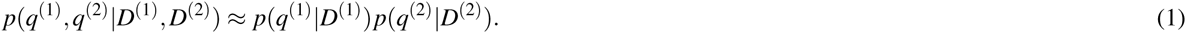

While, in practice, the quality of this approximation will depend upon a number of factors (including the signal-to-noise ratio associated with the individual datasets), it has the advantage of circumventing many of the challenges associated with trying to model *p*(*q*^(1)^, *q*^(2)^ | *D*^(1)^, *D*^(2)^). Moreover, this independence assumption means that we may perform computations for each dataset in parallel, allowing us to scale to potentially large number of datasets (i.e. many more than 2). Having made this independence assumption, we may obtain samples from the joint distribution by sampling independently from each of the factors, *p*(*q*^(1)^*|D*^(1)^) and *p*(*q*^(2)^*|D*^(2)^). For a given fusion function, *f*, we may then consider the set 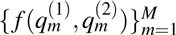, as described above. Extending to *R* datasets is straightforward: we simply obtain samples independently from each *p*(*q*(*r*)*|D*(*r*)) for *r* = 1, *…, R*, and then consider the set 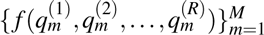.

### GTA: A graph theoretic approach to unsupervised data integration

We start with a collection of datasets, *D*^(1)^, *…, D*^(*R*)^, each of which comprises measurements taken on a common set of *N* entities/items (e.g. genes or patients). We consider the situation in which the prediction, *q*^(*r*)^, associated with dataset *D*^(*r*)^ is the clustering structure possessed by these items. To proceed, we must therefore define: (i) a method for sampling from *p*(*q*^(*r*)^ *D*^(*r*)^); and (ii) a fusion function, *f*, for combining the predicted clustering structures derived from the various datasets. Here, our aim is to identify similarities between the clustering structures associated with each of the datasets.

#### Sampling from p(q^(r)^|D^(r)^)

We consider two different approaches for sampling from *p*(*q*^(*r*)^ *D*^(*r*)^). Where possible, we use Dirichlet process mixture modelling, a nonparametric Bayesian approach, which is discussed in the Supplementary Material. However, in some cases the size of the datasets being considered prohibits the use of such approaches. In these instances, rather than using an ensemble of *M* samples from *p*(*q*^(*r*)^*|D*^(*r*)^), we instead take *M* = 1 and treat the maximum likelihood estimate, 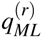, as a single, representative sample.

#### Defining the fusion function

In order to define a fusion function, *f*, we take inspiration from Balasubramanian et al. work12 and adopt a graph theoretic approach. We define **c** to be a clustering of the *N* items of interest (genes, patients, etc.), so that *c_i_* is the cluster label associated with item *i*. We form an *N × N* adjacency matrix as follows:

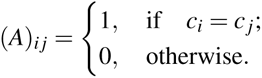

Given a collection of *R* such clusterings, say **c**^(1)^, *…,* **c**^(*R*)^, we may form a corresponding collection of adjacency matrices, *A*^(1)^, *…, A*^(*R*)^, where *A*(*k*) is the adjacency matrix representation of clustering **c**(*k*). The Hadamard (entry-wise) product of these matrices, 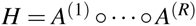, defines a new clustering, in which items *i* and *j* appear in the same cluster if and only if all of the clusterings **c**^(1)^, *…,* **c**^(*R*)^ agree that they should appear in the same cluster. From a graph theoretic perspective, *H* corresponds to the graph formed by taking the intersection of the graphs defined by *A*^(1)^, *…, A*^(*R*)^ (see Figure 1). Assuming that each of the clusterings, **c**^(1)^, *…,* **c**^(*R*)^, corresponds to a different dataset, we define our fusion function *f* to act on these and then return the clustering corresponding to the Hadamard product, *H*. Equipped with this fusion function, and *M* samples from *p*(*q*^(*r*)^ | *D*^(*r*)^) (for *r* = 1, *…, R*), we may proceed to describe with the GTA algorithm (Algorithm 1).

##### Algorithm 1

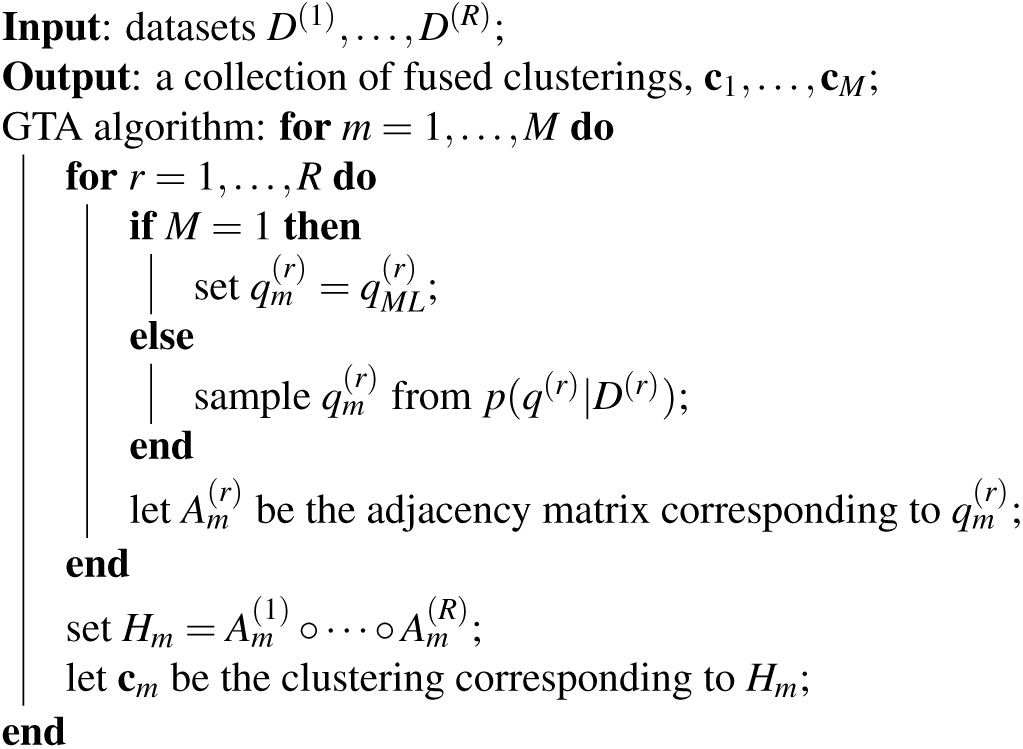

The GTA algorithm.

#### Summarising the fused output

The output of the GTA algorithm is a collection of *M fused clusterings*. While these provide a useful indication of the uncertainty in the fused output, it is often also helpful to condense these into a single, summary fused clustering, 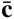. This can be done by constructing a posterior similarity matrix^14^ of 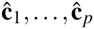 and maximising the posterior expected adjusted Rand index.^15^

#### Heuristic score of clustering similarity

In some instances, it is useful to measure the compatibility between data sources by comparing the independent clusterings (obtained before data integration) with the fused clustering (obtained using the GTA algorithm). Due to the nature of our technique it is expected to observe more clusters *after* the data integration process. This is particularly true when studying less related datasets and when integrating more than a few data sources. We therefore define the following measure of similarity between any two data sources *D_r_* and *D_I_*,

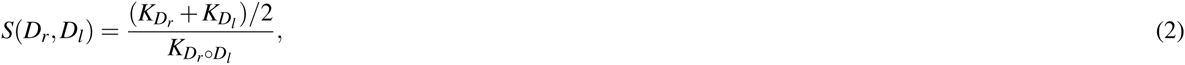

where *K_Dr_* and *K_Dl_* correspond to the number of clusters before data integration, and *K*_*Dr º Dl*_, the number of clusters after integration. Here, the score *S* can have any value between zero and one. The closer *S* is to 0, the more dissimilar datasets are. On the other hand, the closer *S* is to 1, the more substantial the similarity between the structure of the two datasets.

### GTA Applications

In this section we explore the capabilities of GTA to data integration. We start by illustrating the applicability of a heuristic score to identify similarities between several data sources. Then we test GTA performance on popular example applications from the literature, including a dataset for *sporadic Inclusion Body Myositis*.

### Capturing similarities across yeast time courses

We consider a *S. cerevisiae* dataset^16^ that contains *m*RNA transcription levels taken to study the cell cycle. From 416 genes that had previously been identified to have periodic changes in transcript levels we select 100, and assign them into seven clusters using DPM with GPR likelihood model (Figure 2A). The details regarding MCMC specification and diagnostics are summarised in Supplementary Material section B. To demonstrate the performance of GTA, we further consider the artificial dataset example^1^ that was constructed using the same 100 genes. Briefly, this example consists of six data sources, where the first source is the original dataset (see Figure 2**A**), and the other five were obtained sequentially, by randomly permuting a quarter of gene names in the previous dataset. Applying GTA on pairwise combinations of these datasets we identify the numbers of genes that cluster together before and after the fusion. These can be used for computing the score of agreement and for determining the similarities across all 6 datasets. The pairwise similarities are summarised in Figure 2B where columns and rows identify which combination of datasets were considered, and colour illustrates the level of similarity. Alternatively, the similarity between these datasets can be identified from the final clusterings by computing the adjusted Rand index (ARI).^15^ The ARI compares two given partitions of the same list of genes and is based on how often a gene (observation) is associated with the same cluster in both partitions (see Figure 2C).

**Figure 2.**
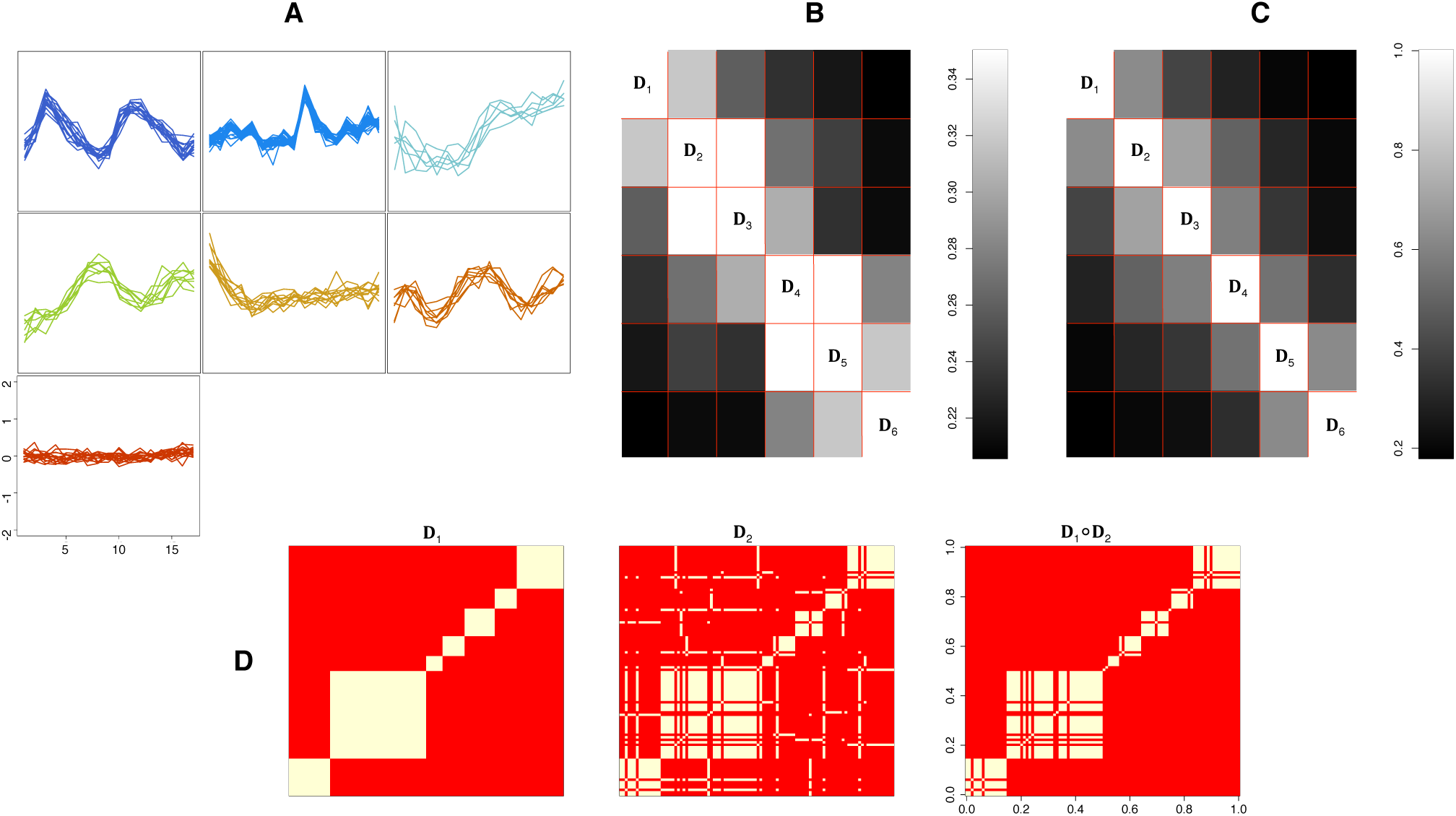
GTA applications to six artificial datasets. **(A)** Illustrates genes from^16^ sorted into seven clusters using DPM model with GP likelihood. **(B)** Illustrates similarity between six final clusterings using similarity measure *S*(*D_r_*, *D_l_*) outlined in ^(2)^. **(C)** Illustrates similarity between the same clusterings using adjusted RAND index. **(D)** Illustrates the effects of Hadamard product – each heatmap corresponds to the adjacency matrix constructed from original dataset, *D*_1_, and the first modified dataset, *D*_2_.

### Integrating cell cycle datasets

We next compare the results from GTA to the results by *Multiple Dataset Integration* (MDI).^1^ We consider integrating two different combinations of datasets from yeast cell cycle studies. The first dataset contains gene expression time courses,^17^ where *m*RNA measurements are taken at 41 time points across 551 genes that exhibit oscillatory expression profiles. The second dataset is ChIP-chip data^18^ that contains binary information about proteins binding to DNA.

Applying the independent DPM models with Gaussian process likelihood to the gene expression time courses, and multinomial likelihood to the transcription factor binding data we construct the adjacency matrices from posterior clusterings. Then, computing the Hadamard products we are able to extract the final allocation variable that contains indices of genes that cluster together in both datasets. As before, further details on MCMC specification and diagnostics are given in Supplementary Material section B.

In this example we are aiming to compare the results from our method to the results from MDI, which jointly clusters all datasets by modelling the dependencies between them. Using MDI approach the final clustering can be extracted by calculating a fusion probability – the probability that any two genes are fused in two datasets, and by removing those genes where this probability is less than 0.5. For this reason, using GTA we can also consider removing genes that lack the evidence of clustering together. This can be achieved by computing the matrix 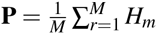, where each matrix entry is probability, *P*_*i j*_, for gene *i* and gene *j* to be in the same cluster in both gene expression and transcription factor binding datasets; and removing genes *i*, where *P*_*i j*_ *<* 0.5 for every *j*. The above procedure is somewhat analogous to MDI in terms of looking only at those genes that are fused across both datasets. Then, applying the Fritsch and Ickstadt^14^ approach to filtered posterior clusterings we obtain the final data partition that assigns genes into clusters based on information from both datasets. Figure 3A,C illustrates the performance of GTA, where genes are allocated into clusters based not only of their expression profiles but also on which transcription factors binds to DNA. This means, GTA enabled us to elucidate those sets of genes that are regulated by the same transcription factors (see supplementary Figure 2, here gene expression time courses on the left are projected on to ChIP-chip dataset on the right). Using GTA we identified 10 clusters that consist of a total of 44 genes (for comparison MDI identified 48 genes). Figure 3A,B illustrates the network structure of yeast cell cycle genes assigned in to clusters using GTA and MDI respectively. Here a line indicates that any two genes are assigned to the same cluster after the fusion. Compared to MDI’s final clustering, GTA in addition identified three clusters whose genes are highly active in G1/S cycle stages and are involved in processes such as: (V) negative regulation of transcription involved in G1/S transition of mitotic cell cycle, regulation of transcription, cellular response to DNA damage stimulus, DNA recombination, DNA repair; (VI) cell wall organisation or biogenesis, lipid transport; (VII) metabolism, regulation of passage through the cell cycle, regulation of cell cycle, cellular response to DNA damage stimulus (see Tables 1 and 2 in sup. mat. for further details).

**Figure 3.**
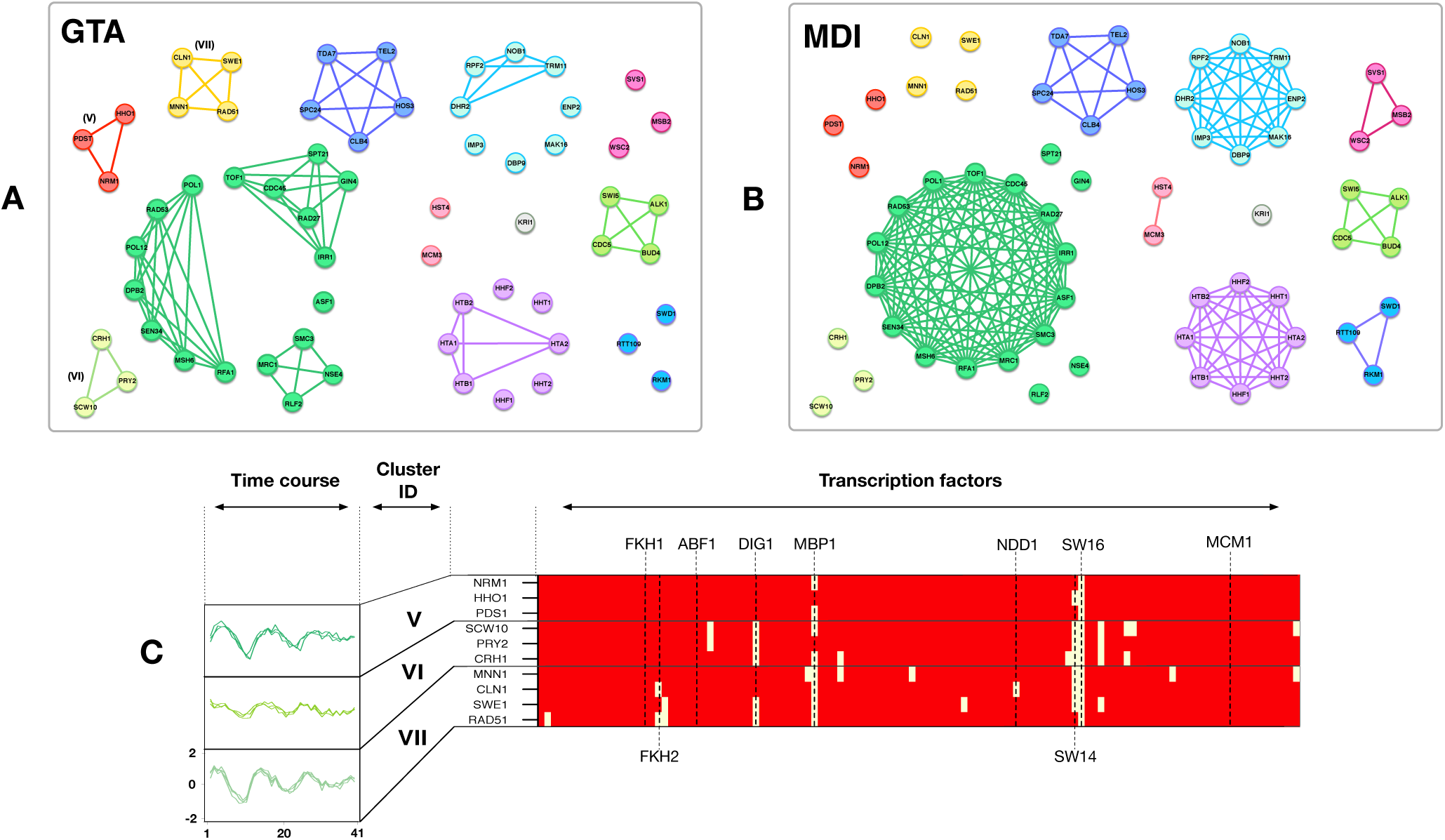
GTA applications to yeast cell cycle datasets. Integrating yeast cell cycle time course and transcription factor (TF) binding datasets. **(A)** Networks illustrates final clustering structure using fused samples from GTA. **(B)** Networks illustrates final clustering structure using MDI. **(C)** Further illustrate 3 additional clusters identified by GTA; on the left – gene expression time courses are projected on a heatmap of TF binding data on the right, here yellow colour indicated that a TF is binding to a gene. Horizontal black lines mark cluster boundaries.

The second step of our yeast cell cycle example considers integrating three datasets: gene expression, transcription factor binding and protein-protein interaction (PPI). In order to also fuse the PPI dataset, we obtain data from BioGRID,^1,19^ and then cluster genes using the DPM model with the multinomial likelihood function. Then, applying GTA we can incorporate the PPI clustering outcome with the gene expression and TF binding results. Again, thinning out the genes that lack evidence for clustering together, we obtain a set of 14 genes that can be assigned into 6 clusters (for comparison MDI assigned 16 genes into 5 clusters). Supplementary Figures 4**A,B** shows final networks fused across three datasets using GTA and MDI respectively; in addition, Supplementary Figure 3 projects clusters from GE data on to TF and PPI. For convenience, Supplementary Material Tables 3 contains further details regarding gene function.

Thus, we can obtain comparable results to MDI without explicitly modelling dependencies between various data sources. This example demonstrates that our data fusion technique is a competitive tool that can be applied for detecting underlying regulatory processes at the molecular level. Moreover, in our case it was not required to rerun all data pre-processing (clustering) in order to additionally consider the fusion of the PPI dataset.

### Breast cancer data

In this example we explore the performance of our data integration technique on a breast cancer dataset, where we aim to integrate four different data sources taken from *The Cancer Genome Atlas*.^20^ We use previously described dataset^2^ that contains of preselected 348 tumour samples measured across four datasets: RNA gene expression (645 genes), DNA methylation (574 probes), *m*iRNA expression (423 *m*iRNAs) and reverse phase protein array (171 proteins). In order to cluster all data sources, we adopt a modified version of Bayesian consensus clustering (BCC) approach.2 BCC is data integration technique that seeks to simultaneously model data specific and shared features by inferring the overall clustering 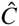 (that describes all datasets) and by inferring data specific clusterings 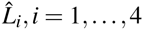. The source specific clustering is controlled by parameter *α* = [*α*_1_, *…, α*_4_], which express the probability of how much each *L_i_* contributes to the overall 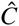. Our goal is to cluster each dataset independently without inferring the overall clustering 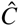. For this reason we fix the probability *α* = 1 and perform BCC individually on each genomic dataset (using publicly available R code). Next, applying GTA on posterior samples *L_i_* we identify the overall clustering 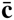.

**Table 1.** Comparison between BCC overall and GTA final clusterings. In the table are given numbers of tumour samples per cluster; e.g. GTA cluster 3 contains 6 samples of *Her2*, 65 samples of *Basal* and 4 samples of *Luminal A*, for this reason, cluster 3 can be summarised as containing mostly *Basal* type tumours. Clusters are classified by particular cancer subtype using publicly available R code.^2^

| TCGA tumor subtypes | BCC single run |  |  | GTA |  |  |
| --- | --- | --- | --- | --- | --- | --- |
|  | Cl<sub>1</sub> | Cl<sub>2</sub> | Cl<sub>3</sub> | Cl<sub>1</sub> | Cl<sub>2</sub> | Cl<sub>3</sub> |
| *Her2* | 20 | 6 | 13 | 5 | 28 | 6 |
| *Basal* | 4 | 2 | **66** | 1 | 6 | **65** |
| *Luminal A* | **81** | **76** | 4 | **59** | **98** | 4 |
| *Luminal B* | **73** | 3 | 0 | **59** | 17 | 0 |

Breast cancer is a heterogeneous disease, for this reason four biologically distinct molecular subtypes where connected to these data sources.^2,20^ They are known as *Her2*, *Basal*, *Luminal A*, *Luminal B*, and are associated with different clinical prognosis.21, 22 In order to assess our results we identify cancer subtypes that are associated with each cluster and compare these to the BCC clusters. Table 1 illustrates the summarised results. In the second column we present the outcome from BCC (a single run of publicly available R code) and in third we summarise results from GTA. It can be seen that clusters identified using our technique can be described by similar cancer subtypes when compared to BCC outcome (e.g. *Cl*_3_ in our case contains mostly *Basal* type tumour samples (65 in total), and this corresponds to *Cl*_3_ in BCC analysis; our *Cl*_2_ can be described by *Luminal A* subtype, and in BCC case *Cl*_2_ is a similar cluster. Furthermore, both cluster, *Cl*_1_, from our method and BCC cluster, *Cl*_1_, contains tumour samples from *Luminal A, B* subtypes).

We compared our method to the BCC final outcome. Although our method does not model the relationships between data specific and overall clustering, the final data integration outcome leads to very similar results.

### Sporadic inclusion body myositis

In this section we apply our technique on clinical gene expression datasets that include: (i) *sporadic Inclusion Body Myositis* (sIBM), which is an inflammatory muscle disease that progresses very slowly, causes muscular weakness and eventually muscle atrophy;^23,24^ (ii) *polymyositis* (PM) which causes chronic inflammation of the muscles; and (iii) a dataset containing human protein-protein interactions (BPPI) (see Supplementary Material, section D for further details). Both sIBM and PM are associated with ageing but interestingly sIBM can be frequently misdiagnosed as PM, and the explicit diagnosis can only be confirmed via a muscle biopsy.^25^ Current understanding is that sIBM is driven by two coexisting processes (autoimmune and degenerative); however, the actions by which sIBM occurs are still only poorly understood.^25,26^

#### PM and sIBM data integration

Here we focus on experimental sIBM and PM datasets that have 5 and 3 data points (clinical cases). In order to apply our technique, we select 424 genes that where previously identified to have the largest variation in their expression across all data points.^27^ In order to cluster these datasets, we employ “*mclust*” package in R, which fits a Gaussian mixture model and uses the Bayesian information criterion to estimate the number of components. Because we do not have access to the clustering samples from the posterior for each dataset, we use GTA with a special case *M* = 1 to integrate single clusterings from both datasets. The integrative analysis enabled partitioning of large PM and sIBM clusters into smaller ones, however this resulted in a greater number of clusters after the fusion, Figure 4 illustrates 12 largest clusters. To validate our results we used a web-based tool called “ToppGene”^2^ that performs gene set enrichment analysis and allows the detection of functional enrichment for phenotype (disease) and GO terms such as biological function. Such analysis enabled us to identify those clusters (5 out of 12) that are enriched with diseases like rheumatoid arthritis and myositis, and biological processes related to response from immune system. This suggests that genes in disease enriched clusters might play an important and shared role in both PM and sIBM, and for this reason they are promising targets for further analysis.

**Figure 4.**
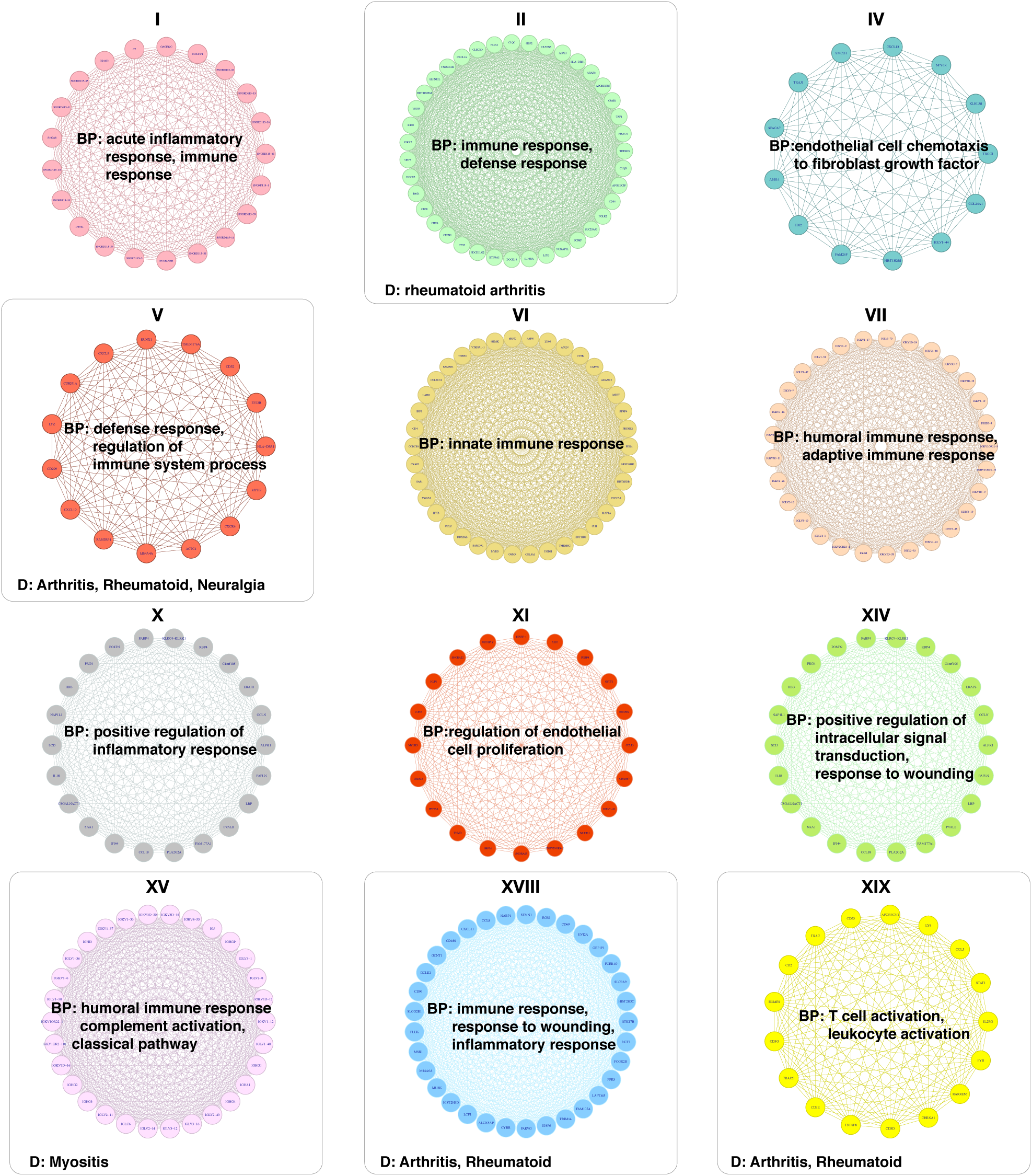
GTA applications to myositis datasets. Illustrate clusters obtained by integrating PM and sIBM gene expression datasets using GTA. Each network represents a fused cluster, which is associated with one of enriched ontologies, biological process (BP), and disease (D).

## Discussion

In this paper we have developed and illustrated an alternative way to integrate data generated from different sources. GTA mainly relies on the outcomes from Bayesian clustering approaches and is based on concepts from graph theory. We have demonstrated that our technique, can produce similar data fusion results when compared to recently proposed data integration methods. Equally, while GTA can fuse the outcomes from Bayesian clustering algorithms, the special case *M* = 1 can be applied in order to fuse a single clusterings across various datasets.

Here we have demonstrated the applicability of our technique to a variety of biological problems: the identification of potentially underlying regulatory mechanisms in the yeast cell cycle, sub-typing tumours in breast cancer data, and exploring similarity patterns across inflammatory muscle diseases. We have compared our technique to MDI and BCC, which are currently used to address similar data integration problems. The main benefits of our graph-theoretical approach include: (i) the applicability to Bayesian and non-Bayesian type clustering approaches. This means that our methodology can be applied in order to model multiple sources on a genome scale data without facing computational challenges; and (ii) ability to perform clustering in a parallel fashion. Such feature might be advantageous in situations where it is necessary to rerun computations in order to consider additional datasets. As part of our modelling approach, we have defined a measure of similarity between two data sources. This can be used to evaluate the effects of data integration routine and in order to assess the agreement between data sources.

We note that it is possible to apply our graph-theoretical approach to the outcomes from simple clustering techniques, for example hierarchical or *K*-means clustering. This could be done by first performing a bootstrapping approach on each gene within a dataset *D_r_*, *M* times; for further details on bootstrapping see^28^ and^29^. This would produce a set of bootstrapped datasets 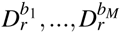. The bootstrapped datasets together with the initial dataset can be clustered using a standard hierarchical clustering algorithm. The clustering outcomes can to some extent be viewed as being the analogues to the samples from posterior. Then after bootstrapping all data sources we can use GTA to perform integrative modelling. This process can be seen as a frequentist modelling approach to data integration.

## Acknowledgements

This work was supported by the Leverhulme Trust (to JŽ and MPHS), the Royal Society (to MPHS), HFSP (to PK and MPHS) and BBSRC (to MPHS)

## Author contributions statement

JŽ carried out the analysis and prepared the manuscript; PK contributed code, participated in the design and coordination of the study and helped to draft the manuscript; MPHS directed the design and coordination of the study and helped to draft the manuscript. All authors read and reviewed the final manuscript.

## Additional information

The authors declare that they have no competing interests.

## Footnotes

1 For the sake of generality, we leave the definition of *q* deliberately vague, but these predictions could be, for example, assessments of disease risk

2 https://toppgene.cchmc.org

